# Mechanistic tradeoffs between local and long-range signaling activity in natural and synthetic morphogens

**DOI:** 10.64898/2026.02.06.704427

**Authors:** Gavin Schlissel, Anders S. Hansen, Pulin Li

## Abstract

Hedgehog family morphogens present an interesting paradox: despite being hydrophobic due to dual-lipid modifications, they form spatial concentration gradients that are highly conserved and essential for many aspects of metazoan development. Using live-cell single-molecule tracking and engineered synthetic signaling ligands, we isolated the distinct contribution of each lipid modification to Hedgehog diffusion and signaling potency. We found that although both lipid modifications enhance signaling potency, they do so through different mechanisms. Palmitate directly promotes receptor engagement, whereas cholesterol topologically confines secreted morphogens on the cell surface, effectively using the lipid membrane as a non-signaling co-receptor that enriches ligands locally at the cost of restricting long-range diffusion. Our results on the function of cholesterol point to an intrinsic tradeoff between signaling potency and gradient formation, with implications for the evolution and mechanism of non-signaling co-receptors.

## Introduction

Signaling proteins form spatial gradients in animal tissues that instruct developmental fate or physiological state, and the shape of signaling gradients can determine the outcome of developmental patterning or immune activation(*1–3*). To understand what factors affect signaling gradient formation, it is critical to understand how ligands travel through the extracellular environment to reach target cells and engage with receptors to trigger signaling responses, because the mechanisms of both ligand movement and signaling potency contribute to the shape of a signaling gradient(*2, 4–7*).

Signaling ligands are often structurally compact and it remains unclear which biochemical features, including their idiosyncratic post-translational modifications, affect ligand movement, signaling potency or both. For example, Hedgehog is modified with two lipids: palmitate on its N-terminus and cholesterol on its C-terminus, and Wnt family ligands are modified internally with a palmitoleate(*8–10*). These hydrophobic modifications are thought to have two major consequences: first, they are thought to reduce ligand solubility in the aqueous extracellular matrix and thus influence the ligand diffusion. Consistent with this idea, Hedgehog cholesterol is known to limit Hedgehog secretion and movement(*11–13*). Second, Hedgehog palmitate and Wnt palmitoleate directly bind their cognate signaling receptors and promote signaling, suggesting that post-translational modifications might impose a tradeoff between diffusivity and signaling potency (*12, 14–16*). Because ligand modifications affect multiple properties of signaling ligands, studies that rely on natural signaling ligands and receptors cannot uncouple the effect of ligand diffusivity from ligand potency during gradient formation.

In addition to intrinsic biochemical properties of ligands, the extracellular environment also plays a central role in shaping signaling gradients by transiently interacting with ligands as they permeate a tissue(*17–19*). For example, cell surface receptors can influence gradient formation by internalizing and degrading ligands as they transduce a signal, and ligand diffusion and degradation rates are both critical in setting the size of a signaling gradient(*4, 5, 20*). In addition to receptors, signaling molecules can also interact with a wide variety of co-receptors that are abundant on cell surfaces. Although co-receptor interactions are canonically understood to modulate ligand/receptor interactions, and thus regulate signaling potency, co-receptors could also affect the extracellular ligand diffusion by transiently binding ligands and modulating their mobility (*21, 22*).

The diverse biochemical features of ligands and the complexity of the extracellular environment highlight the need for isolating each individual interaction to evaluate its effect on diffusion and signaling potency. Our previous work on Hedgehog diffusion revealed that extracellular Hedgehog diffusion is complex and dynamic, involving formation and dissolution of many different biochemical interactions that each last on the order of ~10-300 ms(*13*). Furthermore, the precise on-rates and off-rates that govern the dynamic extracellular interactions are critical for regulating the gradient length scale, and that modulating these rates with the Hedgehog chaperone SCUBE allows the gradient length scale to vary across tissues or organisms. In contrast to bulk assays, including conventional binding assays or techniques like fluorescence recovery after photobleaching (FRAP), single particle tracking captures heterogeneity of diffusion among Hedgehog particles and directly measures the kon and koff of transient Hedgehog complexes, allowing us to measure the relationship between extracellular binding dynamics and gradient formation(*13, 23–26*).

Here, we used single particle tracking microscopy to understand how post-translational modifications to the developmental morphogen Sonic Hedgehog (SHH) regulate its movement through the extracellular environment. We found that the Hedgehog core protein and each Hedgehog lipid modifications had distinct contributions to Hedgehog diffusion dynamics and gradient formation by mediating distinct interactions with Hedgehog co-receptors, lipid membranes and the extracellular matrix. Furthermore, we built synthetic morphogens based on our understanding of Hedgehog biology and found that the morphogen diffusion mechanism is tightly coupled to morphogen signaling potency. This work revealed an underlying biophysical tradeoff between diffusion and receptor engagement, which we speculate could underlie the extensive diversification of co-receptors that has occurred across the evolution of multicellular life.

## Results

### Palmitate and Cholesterol have distinct effects on Hedgehog signaling gradients

Given the high evolutionary conservation of the two lipid modifications on Hedgehog, we set out to understand whether these biochemical features have similar or distinct effects on the Hedgehog gradient formation. We generated clonal sender cell lines that produce alleles of Hedgehog lacking either N-terminal palmitate, C-terminal cholesterol or both lipid modifications and measured their ability to form signaling gradients in a reconstitution assay based on NIH3T3 cells (Fig 1A)(*20*). A HaloTag was inserted in the same internal position E130 of all SHH variants as described previously to enable direct imaging of the ligands(*13, 27*). Fully modified Hedgehog formed signaling gradients that extended ~125-175µm beyond the membrane boundary of sender cells (Fig 1B, C).

**Figure 1.**
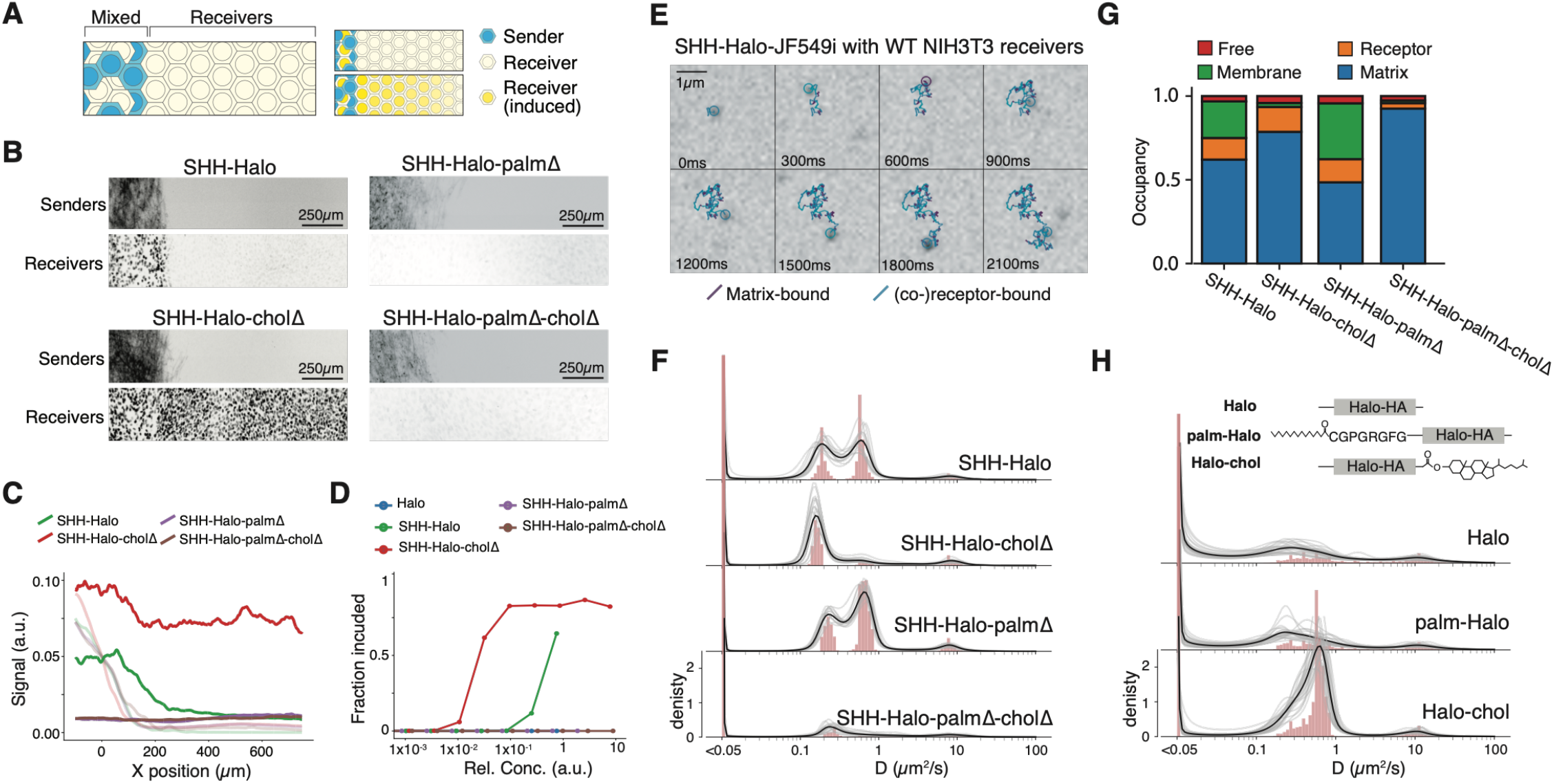
Cholesterol and Palmitate had distinct contributions to ligand diffusion and signaling potency. A. Schematic of signaling gradient experimental design. B. Signaling gradients formed by Hedgehog alleles lacking post-translational modifications. C. Quantification of quasi-1D signaling gradients in (B). X position is measured relative to the boundary of the sender cell zone. Light lines represent sender cell intensity, and dark lines represent receiver cell signal for each condition. D. Signaling activity of SHH variants in NIH3T3 derived SHH reporter cell line cGS401. E. Montage of single-particle trajectory color-coded by instantaneous diffusion rate. F. Histogram of maximum likelihood diffusion rates. Y axis is log-scaled. Gray lines reflect the marginal posterior maximum likelihood of each individual well position. Black line reflects the marginal posterior maximum likelihood summing across all well positions. Histogram bars reflect the frequency histograms of diffusion rates, where each trajectory was assigned a single diffusion rate equal to its maximum likelihood diffusion rate. G. State occupancy of molecules plotted in (F). H. Schematic of HALO-derived alleles and histograms of their diffusion rates.

The different Hedgehog alleles had drastically different gradient phenotypes. A Hedgehog allele with a single amino acid mutation (C24A) that prevented palmitoylation was incapable of signaling, even in the region where sender and receiver cells directly contact each other (Fig 1B, C)(*28*). In contrast, Hedgehog lacking cholesterol due to deletion of the C-terminal Hedgehog auto-processing domain formed long-range signaling gradients over >500 µm(*8*). Because Hedgehog alleles lacking palmitate did not form signaling gradients, it was ambiguous whether palmitate affects the mechanism of ligand movement or simply prevented ligand-receptor interactions. Similarly, the extended signaling range of Hedgehog lacking cholesterol could be explained either by enhanced Hedgehog mobility or changes in Hedgehog potency. Although the two lipids appear to impact Hedgehog signaling gradients differently, whether they have distinct, separable effects on signaling potency and ligand diffusion requires orthogonal assays.

### Cholesterol and Palmitate have distinct contributions to signaling potency and ligand diffusion

To measure signaling potency of different Hedgehog alleles, we generated conditioned media with each Hedgehog allele, then labelled each allele with HTL-JF549i. Because the Halo tag binds HTL-JF549i with 1:1 stoichiometry, we used fluorescence densitometry to precisely measure the concentration of each Hedgehog allele before stimulating receiver cells with the conditioned media. We observed that alleles lacking palmitate could not activate Hedgehog signal in the concentration range tested, whereas Hedgehog lacking cholesterol was more potent than intact Hedgehog (Fig S1A, 1D). This is consistent with the previous findings that the Hedgehog N-terminus, including the N-terminal palmitate modification, binds the core of the Hedgehog receptor Ptch1 during Hedgehog signaling(*15*). In contrast, the enhanced Hedgehog signaling potency by the removal of cholesterol suggests that Hedgehog C-terminal cholesterol limits the ability of Hedgehog to bind Ptch1. Together, these results confirmed that the gradient phenotypes of different Hedgehog mutant alleles are at least partly due to altered signaling potency.

To understand how cholesterol and palmitate affect the diffusion of Hedgehog, we used single-particle tracking to analyze the diffusion of each Hedgehog allele in an extracellular matrix synthesized by wild-type NIH3T3 cells (Fig 1E-G). For all Hedgehog alleles, most molecules bleached in a single step, suggesting that Hedgehog did not form homotypic complexes or micelles (Fig S1B). Consistent with previous results, wild-type Hedgehog diffused in four discrete populations reflecting molecules associated with the insoluble extracellular matrix, with membrane proteins, with the membrane itself, or molecules that were freely diffusing (Fig 1F,G)(*13*).

Tracking the diffusion of different Hedgehog alleles revealed distinct functions of palmitate and cholesterol in regulating ligand diffusion. Specifically, removal of C-terminal cholesterol prevented Hedgehog molecules from diffusing in its membrane-embedded form but had no obvious effect on the population associated with membrane proteins, consistent with our previous observation (Fig 1E-F)(*13*). In contrast, Hedgehog lacking palmitate diffused similarly to wild-type Hedgehog. However, Hedgehog without palmitate and cholesterol had a dramatic decrease in the two populations that interact with membrane proteins or the membrane itself, supporting the notion that lipid modifications mediate Hedgehog interactions with diverse binding partners.

Experiments based on natural Hedgehog inherently conflate interactions that depend on palmitate or cholesterol with interactions that depend on the Hedgehog core protein. To isolate the effect of lipid modification, we applied Hedgehog lipid modifications to a simple Halo protein, to test how each modification influences Halo diffusion. Modified Halo proteins were efficiently secreted and bleached in a single step, suggesting they diffused as monomers (Fig S1C). We found that Halo and palmitoylated Halo did not interact with cell surfaces, however cholesterol-modified Halo diffused at ~0.9 µm2/s, similar to the diffusion rate of membrane-embedded Hedgehog molecules (Fig 1H, S1A).

Together, these results revealed the non-interchangeable role of the two lipids in confining Hedgehog to the cell surface. Specifically, cholesterol modification was necessary and sufficient for tethering Hedgehog to the lipid membrane, diffusing at ~0.9 µm2/s. In contrast, both lipids were involved in interaction with membrane proteins that diffused at ~0.2 µm2/s. The fact that removing both lipids is required for eliminating the ~0.2 µm2/s population suggests that the two lipids either modulate the interaction of Hedgehog with non-overlapping binding partners, or both lipids strengthen the interaction of Hedgehog with the same binding partner. In either case, the lipids alone are not sufficient for such interaction; instead, they modify the binding between core Hedgehog protein and the binding partner(s).

### Slow single-particle tracking revealed matrix-associated Hedgehog

Tracking Hedgehog or Halo ligands consistently identified many ligands that were immobile in the cell culture, consistent with interactions between ligand and the extracellular matrix. To determine if these extracellular matrix interactions were a generic feature of any secreted ligand, or a biological feature of Hedgehog that reflects a specific biochemical binding interaction, we modified our single-molecule microscopy approach to observe the slowest Hedgehog molecules. In fast SPT experiments, we use high laser power and fast shutter speeds to measure the instantaneous diffusion rate of extracellular ligands. In contrast, to image immobile ligand we image with very long shutter speeds under very low laser power to limit photobleaching, causing mobile molecules blur into the background (Fig 2A)(*29–32*). This approach allowed us to measure the amount of time that each ligand remains in its immobile state.

**Figure 2.**
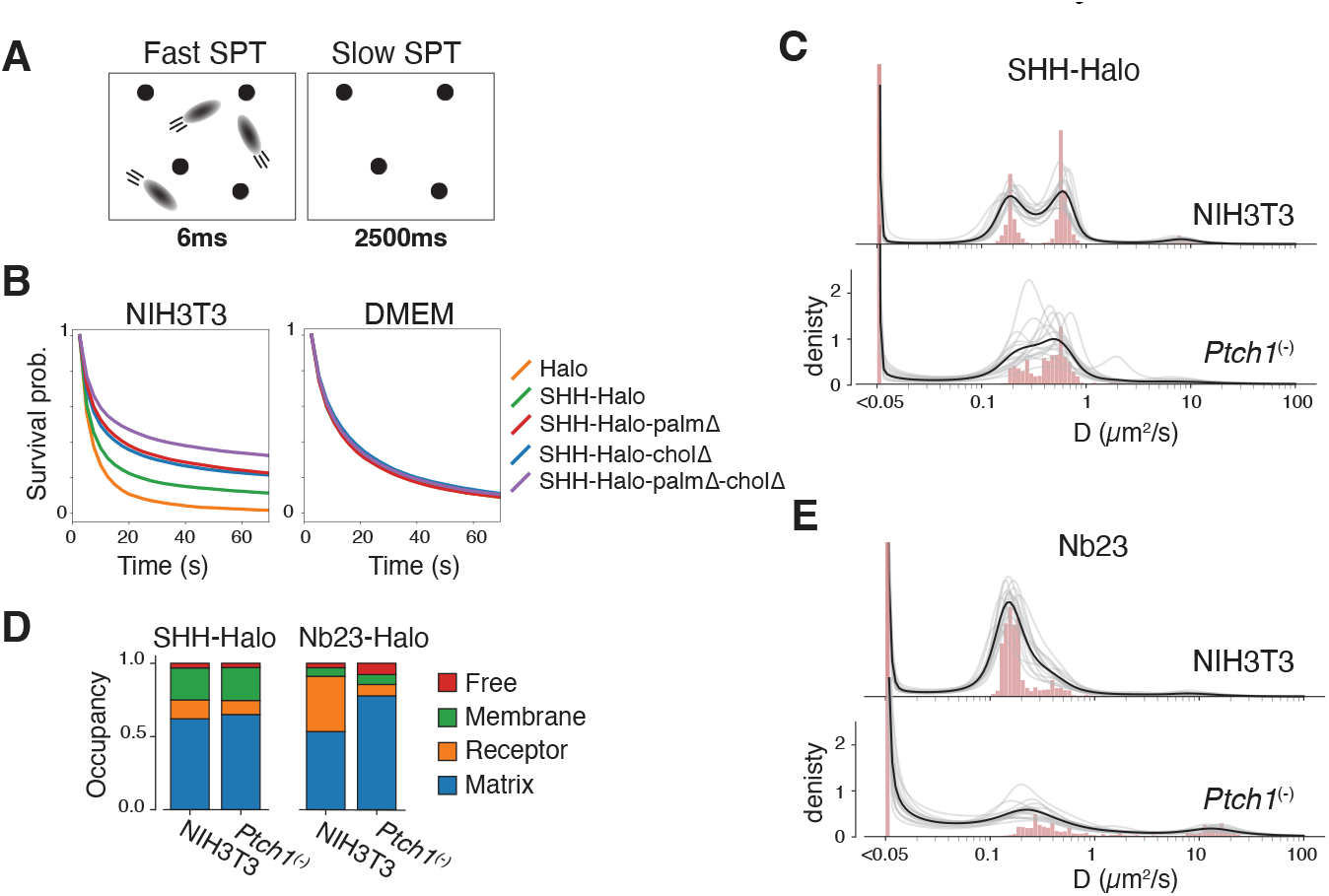
The two slowest populations were associated with immobile matrix and receptor-like molecules. A. Schematic illustrating the design of Slow SPT experiments. B. Survival probability of immobile molecules measured by Slow SPT in a cultured cell extracellular matrix (left) or of purified molecules applied to a cell culture well in DMEM (left). C. Histogram of diffusion rates measured by fast SPT for SHH in the presence or absence of the Ptch1 receptor. D. Quantification of state occupancy for ligands tracked in (C) and (E). E. Histogram of diffusion rates measured by fast SPT for Nb23 in the presence or absence of the Ptch1 receptor.

Using slow SPT, we first found that Hedgehog molecules entered the immobile fraction in a biologically meaningful way. Notably, whereas alleles of Hedgehog showed distinct abilities to bind the extracellular matrix deposited by live cells, purified ligands showed identical interactions with the glass cell culture substrate (Fig 2B, S2A). Furthermore, Hedgehog remained in its immobile state longer than the Halo protein when sender cells were cultured with live receiver cells, suggesting that Hedgehog interacts specifically with certain features of the extracellular matrix (Fig 2B). As with mobile molecules, immobile molecules generally bleached in a single step, suggesting they did not form homotypic complexes in their immobile state (Fig S2B). Notably, a small number of Halo and SHH molecules showed multi-step bleaching when purified ligands were imaged on the glass cell culture substrate, which likely reflects the presence of misfolded or aggregated protein in the protein purification (Fig S2B). Interestingly, hemi-modified and unmodified SHH alleles interact more durably with the extracellular matrix than the dually lipid modified SHH (Fig 2B).

### Membrane-bound non-receptor proteins contribute to receptor-like diffusion

Because Hedgehog N-terminal palmitate was required for Hedgehog signaling through its canonical receptor Ptch1, we were surprised to find that Hedgehog lacking palmitate diffused at a rate that represents association with membrane proteins, among which Ptch1 is the most obvious candidate. We wondered whether the population of Hedgehog molecules diffusing at ~0.2 µm2/s reflected an authentic ligand-receptor interaction. Therefore, we generated receiver cell lines with frameshift mutations in Ptch1 and found that in the absence of Ptch1, Hedgehog nonetheless diffused at a rate that reflected interaction with membrane proteins, with a very minor reduction in the fraction of molecules apparently interacting with membrane proteins (Fig 2C,D).

To validate our Ptch1 mutant cell line and to determine if the Hedgehog molecules that diffuse at ~0.2 µm2/s truly include Hedgehog-Ptch1 complexes, we developed a novel reagent, in which we fused a Ptch1-specific nanobody Nb23 to the Halo tag, and tracked its extracellular diffusion(*33*). We found that the Nb23 synthetic ligand diffused in three populations, with molecules that were immobile, molecules that diffused at ~0.2 µm2/s, and molecules that diffused at ~10 µm2/s (Fig 2E). In contrast, when the Nb23 sender cells were co-cultured with Ptch1(-) receiver cells, Nb23 did not diffuse in a population at ~0.2 µm2/s (Fig 2D,E). Taken together, this suggests that the population of Hedgehog molecules diffusing at ~0.2 µm2/s could reflect both canonical Shh-Ptch1 interactions in addition to interactions with other membrane-bound proteins.

### Each receptor-like molecule has a small contribution to Hedgehog interactions

Because synthetic ligand Nb23 strictly relies on Ptch1 to diffuse at ~0.2 µm2/s, whereas Ptch1 only contribute marginally to the ~0.2 µm2/s population for wild-type Hedgehog, we speculated that the mobile Hedgehog molecules diffusing at ~0.2 µm2/s included interactions with receptor-like molecules that confined Hedgehog. Prior characterization of Hedgehog signal transduction has identified several factors that affect Hedgehog signaling, including the canonical Patched receptor (Ptch1), and the co-receptors Cdon, Boc and Gas1(*34*). Additionally, the GPI-anchored protein family members Glypican 1, 4 and 6, which are homologous to the fly protein Dally-like protein (Dlp), are thought to bind the Wnt palmitoleate, leading to speculation that Glypicans can similarly bind the Hedgehog palmitate and mediate its solubility(*35–38*). Importantly, all of the above-mentioned proteins are expressed in NIH3T3 cells and could contribute to the diffusion dynamics we have previously observed (Fig S3A).

To determine the contribution of each protein to Hedgehog confinement in its receptor-like population, we developed NIH3T3 clones with mutations in Cdon, Boc, Gas1, or Gpc1/4/6. These clones reflected either complete knockout (Boc, Gas1, Gpc6) or ≥ 50% knockdown (Cdon, Gpc1, Gpc4) of the corresponding genes (Fig S3B). Similar to Ptch1 knockout, we did not observe significant reduction in the wild-type Hedgehog population that diffused at a rate consistent with a membrane-embedded protein interaction when any of the membrane proteins were perturbed (Fig 3A). In addition to interactions with the Dlp homologs Gpc1/4/6, Hedgehog has been found to bind Dally homologs Gpc3/5; however, Gpc3/5 were not expressed in our mouse NIH3T3 cells (Fig S3A)(*39*). Therefore, we did not consider their contribution to Hedgehog diffusion in NIH3T3 culture. 37

**Figure 3.**
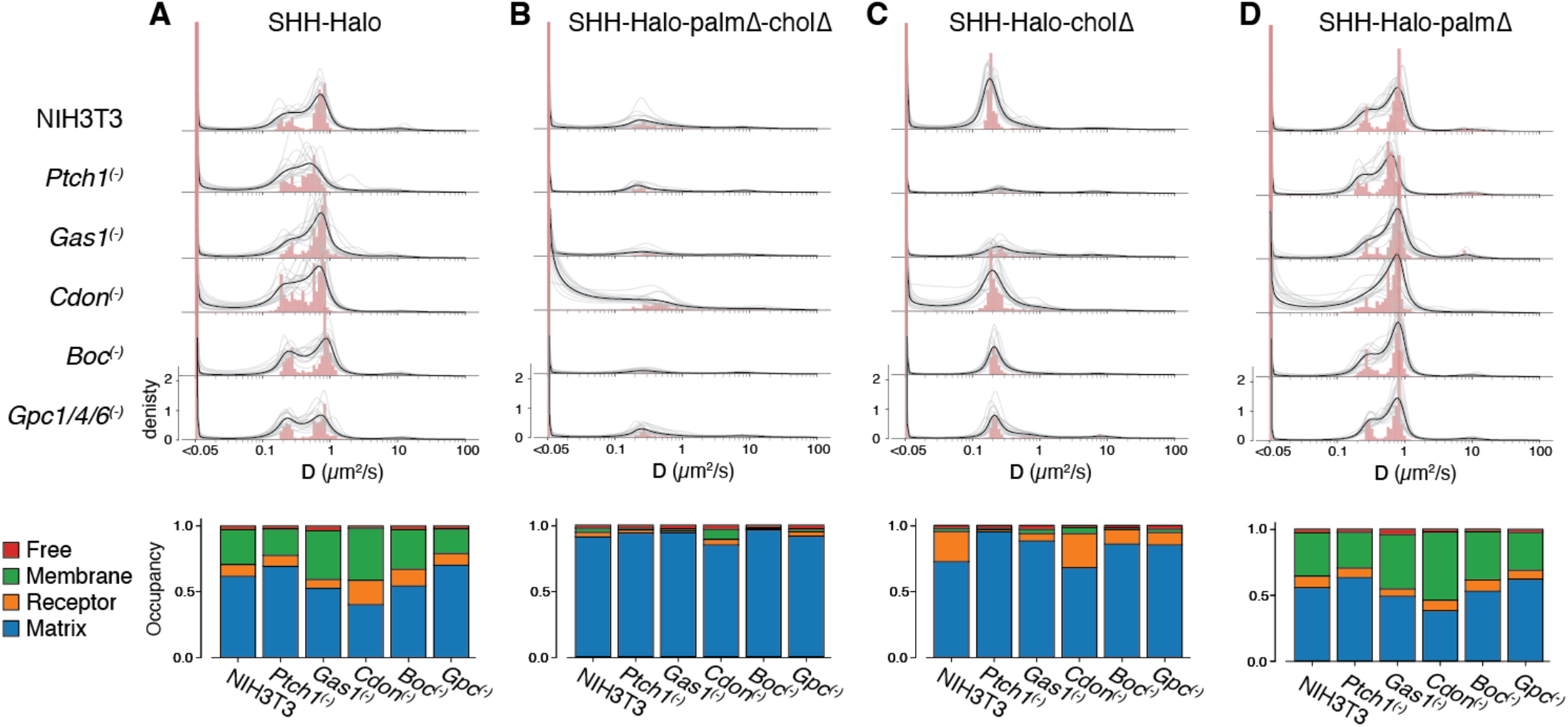
Cholesterol and palmitate interacted with different receptor-like molecules. A. Histogram of diffusion rates for wildtype Hedgehog with mutant NIH3T3-derived receiver lines (top) and quantification of occupancy of each biochemical state (bottom). B. As in (A), but tracking SHH-palmΔ-cholΔ. C. As in (A), but tracking SHH-cholΔ. D. As in (A), but tracking SHH-palmΔ

In addition to using NIH3T3-derived clonal knockout cell lines, we obtained mouse embryonic fibroblasts (MEFs) derived from mice with germline mutations in co-receptors Boc and Cdon(*34, 40*). In both primary cell cultures, Hedgehog diffused in four identifiable populations, similar to its diffusion in NIH3T3 cells. Hedgehog distribution among the four populations in Boc-/-MEF closely resembled Boc-/-NIH3T3 cells, whereas a more severe loss of the receptor-like population was observed in Cdon-/-MEFs compared to NIH3T3-derived Cdon mutant cells (Fig 3A, S5C,D). The relative contribution of each co-receptor depends on the abundance of all co-receptors. Because MEFs are not clonal, it is difficult to know if the expression levels of co-receptors in MEFs are directly comparable NIH3T3 cells, however the results from primary cells qualitatively resembled the results obtained from immortal clonal cell lines.

### Each Hedgehog modification facilitated interaction with distinct sets of receptor and co-receptors

Next, to ask whether Hedgehog lipid modifications modulate the interaction of Hedgehog with Ptch1 and co-receptor proteins, we repeated single-molecule imaging experiments using sender cells that secreted mutant forms of Hedgehog lacking one or both hydrophobic modifications. We found that the Hedgehog core protein interacted weakly with receptor and co-receptor molecules, as shown by the drastically reduced receptor-like population and membrane populations compared to the wild-type Hedgehog (Fig 3B). These rare populations were not affected by mutation to Ptch1 or any of the co-receptors, suggesting that either or both lipid modifications are essential for Hedgehog interactions with its receptor and co-receptors. Indeed, when palmitate or cholesterol was added to the Hedgehog core protein, the receptor-like population re-appeared (Fig 3C, D).

Furthermore, palmitate and cholesterol mediated distinct interactions with the receptor and co-receptors. In the presence of only palmitate, removal of Ptch1 or Gas1, but not other proteins, eliminated the receptor-like population, suggesting that palmitate primarily contributes to interaction with Ptch1 and Gas1 (Fig 3C). In contrast, in the presence of only C-terminal cholesterol, but no N-terminal palmitate, Hedgehog diffusion was not affected by Ptch1, Gas1, Cdon, Boc or Gpc1/4/6 knockout (Fig 3D). We speculate that the cholesterol-modified Hedgehog core protein is efficiently embedded in cell membranes, allowing Hedgehog to interact with receptor-like molecules even in the absence of the N-terminal palmitate interaction surface. In contrast, when Hedgehog is only modified by palmitate it did not embed efficiently in membranes, and interactions between Hedgehog and receptor-like molecules were only detected if they were very high-affinity. Together, these results point to non-overlapping roles of the two hydrophobic modifications in regulating the interaction between Hedgehog and various receptor-like proteins on the cell surface. Consistent with this model, whereas SHH-Halo-cholΔ senders formed uniform signaling fields when cultured with wild-type receiver cells, SHH-Halo-cholΔ formed weak, short-range gradients in receiver cells lacking Gas1 suggesting that Gas1 augmented signaling when SHH-Halo was not capable of cholesterol-mediated membrane-associated diffusion (Fig S3E).

Prior work on Hedgehog co-receptors suggests that each co-receptor is individually not required for Hedgehog signaling, and that embryos with mutations in either Cdon, Boc or Gas1 form normal Hedgehog gradients in their neural tube(*34*). Our data are consistent with the model that each co-receptor has a small individual contribution to promoting Hedgehog-Ptch1 interactions. Furthermore, whereas prior work has found that the co-receptors Cdon, Boc and Gas1 form a relay to strip Hedgehog from its diffusion chaperone Scube, we found that Gas1 additionally contributed to Hedgehog-receptor interactions in the absence of Scube, as NIH3T3 cells do not produce Scube naturally(*41*).

### Cholesterol modification confined Nb23 diffusion

Based on our findings that cholesterol modulated Hedgehog diffusion, and that palmitate was necessary for Hedgehog-Ptch1 interactions, we wondered whether these lipid modifications could be leveraged to manipulate the behavior of the synthetic morphogen derived from Nb23. We built Nb23 alleles that included either palmitate, cholesterol or both (Fig 4A). Because palmitate modification requires a short peptide sequence on the N-terminus of Hedgehog after signal sequence cleavage (CGPGRGFG), as a control we also created Nb23 alleles with the same peptide sequence containing a C24A mutation (AGPGRGFG) to prevent palmitoylation (Fig 4A). For all Nb23 alleles, most molecules (>~95%) bleached in a single step suggesting they diffused extracellularly as monomers, although a small fraction of modified Nb23 molecules bleached in multiple steps, suggesting that the hydrophobic modifications might partially cause Nb23 proteins to oligomerize (Fig S4A).

**Figure 4.**
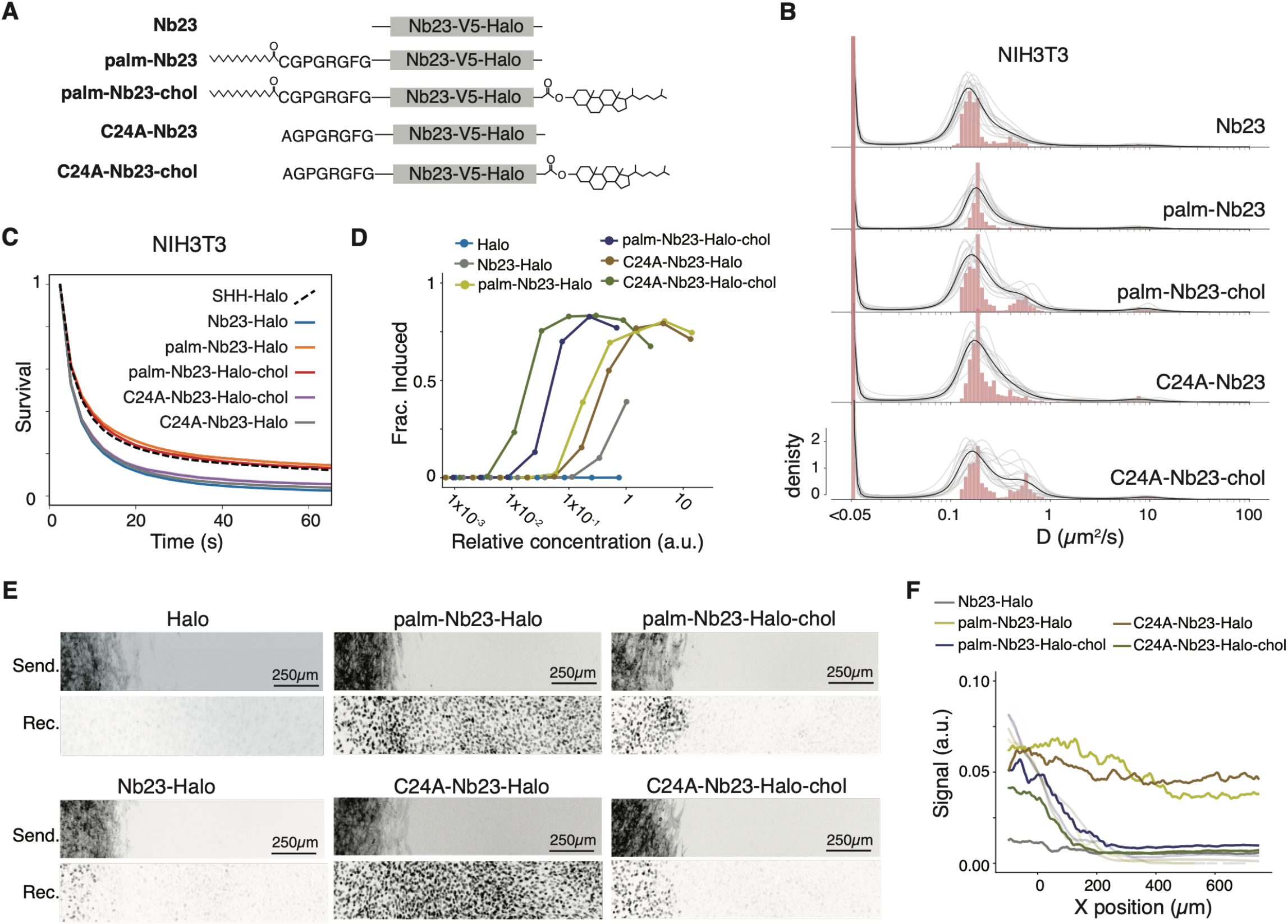
Palmitate and Cholesterol modulated diffusion and potency of a synthetic signaling ligand. A. Schematic of Nb23-derived synthetic morphogens. B. Histogram of diffusion rates for Nb23 and Nb23-derived alleles with wild-type NIH3T3 receiver cells. C. Survival probability of immobile molecules measured by slow SPT in an extracellular matrix synthesized by NIH3T3 cells. D. Specific signaling activity of each Nb23 allele, calculated as the fraction of receiver cells induced by each Nb23 allele. E. Signaling gradients formed by each Nb23 allele. Senders and receivers were plated as in Fig 1A. F. Quantification of signaling gradients in (E). X position is measured relative to the boundary of the sender cell zone.

As expected, modified Nb23 alleles had distinct diffusion features. All alleles diffused as a ~0.2 µm^2^/s receptor-like population that strictly depended on Ptch1, suggesting that Ptch1 is the only membrane protein that these Nb23 alleles bind (Fig 4B, S4B). Alleles containing a C-terminal cholesterol also diffused at ~0.9 µm^2^/s, similar to the diffusion rate of membrane-confined Hedgehog (Fig 4B). This membrane-associated diffusion was independent of Ptch1 (Fig S4A). Interestingly, the cholesterol-modified Nb23 was biased towards interacting with Ptch1 rather than the cell membrane, unlike Hedgehog which is roughly balanced between receptor-like interactions and membrane interactions (Fig 4B, Fig 1E). This suggests that Nb23 bound the Ptch1 receptor with higher affinity than Hedgehog, resulting in an equilibrium state occupancy that favors the receptor-bound form over the membrane-embedded form. Palmitate-modified Nb23 diffused similarly to unmodified Nb23, without embedding in membranes; however, Nb23 that was modified with only palmitate showed a slight increase in matrix-associated molecules compared to unmodified or dually-lipidated Nb23 alleles (Fig 4B, S4B-C).

Because the palmitoylation motif caused only a minor shift in Nb23 diffusion, we used an additional assay to infer whether the palmitoylation motif was successfully modified in culture. Briefly, we speculated that if the Nb23 allele is properly modified, it might interact with the lipid-binding protein albumin(*42*). Therefore, we precipitated unlabeled Nb23, Palm-Nb23 and C24A-Nb23 from conditioned media and used SlowSPT to measure the timescale of Nb interactions either with naked glass or with albumin-coated glass. Each protein interacted similarly with naked glass, but only palmitoylated proteins interacted with albumin-coated glass, suggesting that palmitoylation was intact in Palm-Nb23 S4D,E).

In addition to measuring their instantaneous diffusion rate, we measured the survival of Nb23 molecules in the matrix-associated population using slow SPT. We found that palmitoylated Nb23 ligands bound the matrix to a similar degree as fully modified Hedgehog, whereas Nb23 ligands lacking palmitate showed less stable matrix interactions (Fig 4C). This suggests that the immobile extracellular matrix included some factor capable of binding palmitate, however we have not identified the specific factor.

### Both cholesterol and palmitate enhance Nb23 signaling potency

Prior work on the Nb23 nanobody revealed that Nb23 could activate Hedgehog signaling by trapping the active configuration of the Ptch1 receptor(*33*). Having seen that Hedgehog post-translational modifications could modulate Nb23 diffusion, we next asked whether these modifications could also modulate Nb23 signaling potency. We generated conditioned media containing each Nb23 variant and stimulated Hedgehog receiver cells with each variant (Fig S5A). To our surprise, both cholesterol modification and the Hedgehog N-terminal-derived peptide augmented Nb23 signaling potency, either in its palmitoylated state or when it was mutated (C24A) to prevent palmitoylation (Fig 4D).

Interestingly, the enhanced signaling potency was additive between the N-terminal peptide and cholesterol modification, suggesting that they act through a different molecular mechanism, which is consistent with the finding that the Hedgehog C-terminal cholesterol does not directly access a similar surface of Ptch1 as does the N-terminal palmitate(*15, 43*). It was previously reported that the 8 amino-acid long N-terminal palmitoylation motif (CGPGRGFG), which immediately follows the Hedgehog secretion signal, makes extensive contact with the Ptch1 receptor(*15*). This suggests this peptide motif can participate in a cooperative binding reaction when tethered to Ptch1 by Nb23 and directly enhance signaling, possibly through a previously described “pincer” mechanism, in which the N-terminal peptide and the C-terminal cholesterol pinch the Ptch1 receptor from two sides to activate signaling, although either component of this pincer mechanism was apparently able to enhance Nb23 signaling independently of the intact pincer(*43*).

Notably, the contribution of palmitate and the palmitoylation motif to Ptch1 inhibition was contingent on the proteins that they are appended to. The palmitoylated motif alone was not sufficient to stimulate receiver cells when attached to the free Halo tag, but it was required for Hedgehog to stimulate Ptch1 (Fig S5B,C). In contrast, the Hedgehog N-terminal peptide was sufficient to enhance Nb23 signaling activity, regardless whether it was palmitoylated or not. This effect, in which a weakly binding peptide is converted into a strong agonist by local tethering to a receptor, has been recently described for synthetic GPCR agonists, and likely reflects a generalizable engineering strategy to use cooperativity to improve the potency of synthetic signaling ligands(*44*).

### The two lipids differentially shape signaling gradients formed by synthetic morphogens

Given the distinct effects of each Nb23-derived synthetic ligands on signaling activity and diffusion, we tested how these modifications impact the capability of these ligands to form signaling gradients like a natural morphogen. We found that the unmodified Nb23 could induce receiver cells weakly in their immediate vicinity, but no activation was observed over a long range, consistent with the finding that the unmodified Nb23 is a weak signaling agonist (Fig 4E-F). In contrast, Nb23 alleles containing the N-terminal palmitoylation motif signaled efficiently at a distance, and uniformly activated receiver cells, forming a signaling gradient with a very long (>300 µm) lengthscale (Fig 4E-F). This aligns with the observation that this motif makes Nb23 into a more potent ligand but does not constrain its diffusion, even when the motif is palmitoylated.

Lastly, cholesterol-modified ligands—which had the highest specific potency of all the signaling ligands— formed step-like signaling gradients, efficiently inducing receiver cells in the direct vicinity of the sender cells but not inducing cells away from the source (Fig 4E,F). In principle, shorter signaling gradients could reflect either increased ligand degradation (i.e. through receptor-mediated endocytosis) or decreased ligand diffusion(*5*). Both mechanisms could contribute to shortening the signaling gradients generated by Nb23 alleles tagged with cholesterol. However, we speculate that the gradient shape qualitatively becomes discrete and step-like because cholesterol embedded Nb23 in cell membranes, preventing efficient diffusion through the tissue culture media or extracellular matrix. This interpretation is consistent with our observation that cholesterol is sufficient to embed ligands in topologically defined cell membranes, and with our prior theoretical description of the influence of cell topology on gradient formation (Figs 1H, 4B, S4A)(*13*).

We speculate that the cholesterol modification enhanced the signaling potency of the synthetic Nb23 by confining Nb23 to cell membranes, allowing Nb23 to search for its receptor in the 2D plane of the membrane rather than in the 3D space of the cell culture. In the case of Nb23-cholesterol and in the case of cholesterol-modified Hedgehog, the ligand-membrane interaction is so strong that it severely biases the ligand towards a local 2-dimensional receptor search while limiting long-range gradient formation, revealing an intrinsic tradeoff between the efficiency of a local ligand-receptor search and the ability of a ligand to form a long-range signaling gradient. Under this model, ligands that are highly partitioned into cell membranes can signal efficiently, but only at short range, whereas for ligands that are poorly partitioned into membranes, signaling depends directly on their affinity for their receptor (Fig 5A,B). Importantly, this framework is not mutually exclusive with other mechanisms that regulate the lengthscale of a signaling gradient: for a given amount of membrane partitioning, the signaling potency can further determine the gradient lengthscale by modulating the degradation rate of the ligand(*5*). Instead, we propose that ligand partitioning reflects a separate evolvable property of signaling ligands that can refine gradient shapes without requiring evolution to explore variation in core ligand-receptor interactions.

**Figure 5.**
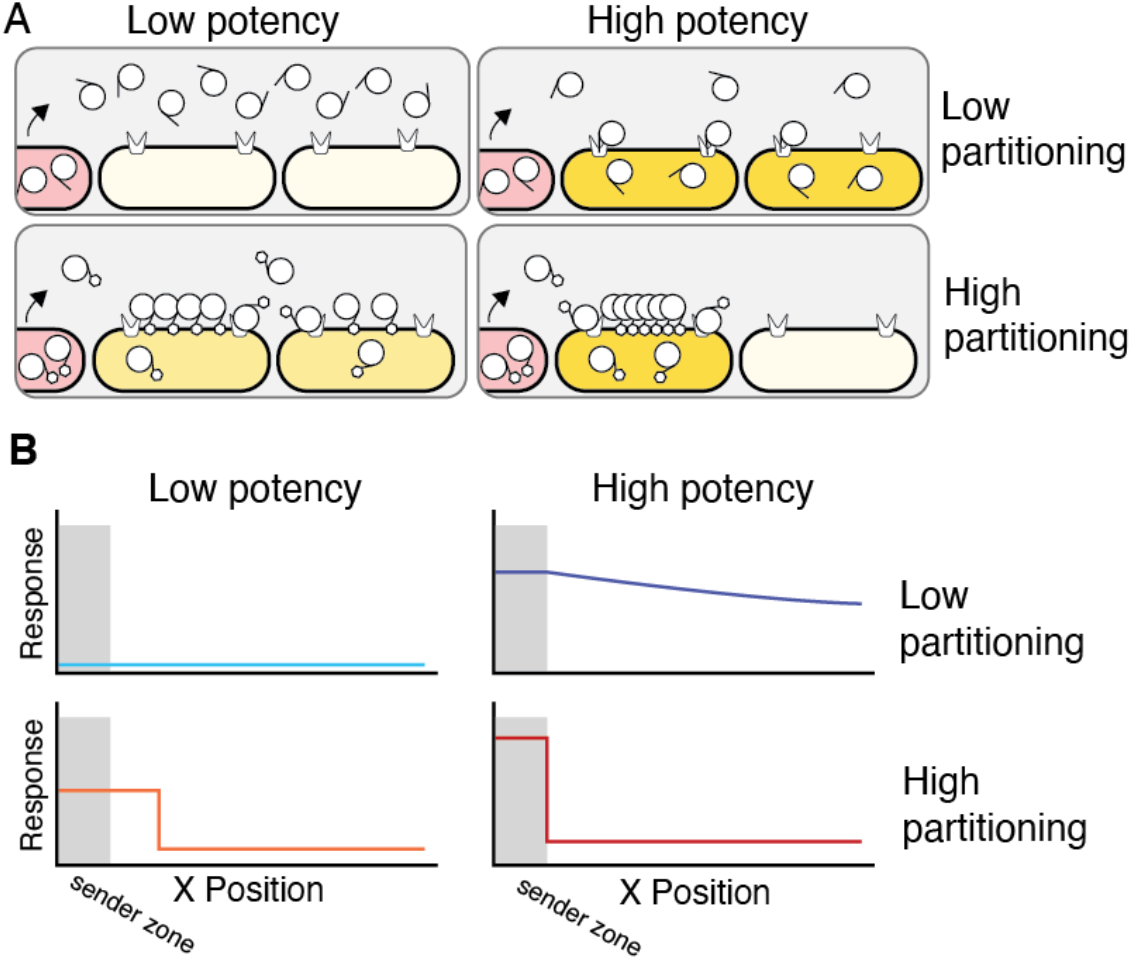
Conceptual model of ligand partitioning and gradient formation. A. Schematic representation of ligand movement under possible combinations of ligand potency and partitioning. B. signaling gradients formed under assumptions in (A). ligand partitioning and ligand potency combine to determine the lengthscale and intensity of signaling gradients.

## Discussion

Hedgehog family morphogens represent a biochemical paradox: despite the fact that the mature Hedgehog signaling ligand is extremely hydrophobic, it forms signaling gradients in tissues by diffusing through hydrophilic environments. This paradox has led to persistent speculation about the mechanism of Hedgehog diffusion, and persistent disagreement about how the Hedgehog N-terminal palmitoylation and C-terminal cholesterol modification affect Hedgehog signaling activity and gradient formation(*43*). Using live-cell single particle tracking, we resolved several long-standing questions about how Hedgehog post-translational modifications affect its diffusion and receptor engagement *in situ*, and we established methods that will be generally useful to understand the biology and evolution of morphogen gradient formation.

Notably, we found that the two Hedgehog lipid modifications interacted differently with annotated receptors, co-receptors and membrane lipids, leading to their differential contribution to signaling potency and ligand diffusion. Although we originally expected Hedgehog palmitate and cholesterol to both contribute to membrane tethering, we found that palmitate was not sufficient to localize Hedgehog to membranes, consistent with prior work in the KRAS field that has found that single fatty modifications are not sufficient to localize KRAS to the membrane without participation of additional chemical factors(*45*). Although Hedgehog palmitoylation did not recruit Hedgehog to cell membranes, we found that palmitoylation together with the 8 amino acids on the Hedgehog N-terminus interacted with Ptch1 and Gas1, directly enhancing receptor engagement. In contrast, cholesterol-modification did not affect Hedgehog-receptor interactions observed by single-particle tracking and Hedgehog alleles lacking cholesterol were more potent than fully modified alleles, despite prior reports that Hedgehog lipid modifications simultaneously engage two surfaces of the Ptch1 receptor(*43*). Interestingly, in mice, Hedgehog lacking cholesterol signals weakly over short distances in the neural tube but can signal over a longer distance compared to wild-type Hedgehog(*12*). This was previously interpreted to mean that Hedgehog lacking cholesterol was hypomorphic for signaling: instead, in light of our observations, we interpret this result to mean that Hedgehog lacking cholesterol diffuses rapidly and permeates the entire neural tube, resulting in more net signaling at a distance and less net signaling near the source compared to wild-type Hedgehog. Importantly, our experiments utilized a tagged allele of Hedgehog that included a large Halo tag insertion at position E130. Therefore, although we could infer a biochemical interaction between the tagged Hedgehog allele and Ptch and Gas1, we cannot rule out the possibility that the Hedgehog-E130 HaloTag prevented identification of interactions between Hedgehog and the coreceptor Cdon or Boc, or Gpc1,4,6.

The tradeoff between signaling potency and long-range diffusion likely reflects a general design principle for morphogens beyond Hedgehog. Despite the lack of cholesterol modification among other morphogens, interaction with co-receptors can partition other morphogens into the plane of the membrane without transducing a signal(*7, 21, 46*). The overall consequence of high membrane partitioning is enhanced signaling closer to the source at the cost of reduced spreading over longer distance(*12*) (Fig 5). In fact, recently developed GFP-derived synthetic morphogens have been found to signal more potently and over longer distances when receiver cells express non-signaling co-receptors to partially confine GFP enabling it to search for its receptor by diffusing in 2-dimensions along the plane of a cell membrane(*21*). We speculate that evolution might similarly use binding interactions between ligands and non-signaling co-receptors to allow organism- or tissue-specific membrane-confined diffusion mechanisms that could modulate both ligand activity and long-range pattern formation. Furthermore, we speculate that design of therapeutic proteins could be improved by bivalent strategies that rationally engage co-receptors or the membrane itself to enhance ligand/receptor interactions.

Remarkably, many diverse biological search problems have evolved to use a similar dimensionally reduced search mechanism to accelerate protein-protein encounters(*47–50*). In the case of receptor tyrosine kinase signaling, membrane-tethering is known to accelerate the interaction between Sos and Ras, resulting in enhanced signal transduction (*47*). In the case of transcription factors searching for their cognate promoter motifs, transient tethering and 1-dimensional “sliding” along chromatin surfaces allows low-abundance transcription factors to find their low-abundance targets efficiently, which would be intractable if transcription factors were limited to searching in 3 dimensions(*48–50*). We speculate that dimension reduction is a deeply embedded feature of biological targeting problems that likely evolved convergently in many unrelated search-limited biological processes.

## Materials & Methods

### Cell culture

NIH3T3 cells were cultured in DMEM (Gibco) with 1mM pyruvate (Gibco), 10% HyClone Cosmic Calf Serum (Cytiva) and 100/mL Penn/Strep (Gibco). Cells were passaged 1:10 at confluence. MEFs were a gift from Benjamin Allen and were cultured as NIH3T3 cells except with 10% Tet-approved Fetal Bovine Serum (Takara) in lieu of Cosmic Calf Serum. MEFs were immortalized by serial passaging at 1:5 dilution factor. SHH receiver cells were designed as described previously, and sender cells were induced to secrete SHH or a synthetic signaling ligand using 250nM 4-OHT as described previously(*13*).

### Generation of synthetic ligands

Throughout the manuscript, proteins that were palmitoylated included a signal sequence and palmitoylation motif derived from mouse Sonic Hedgehog, with the sequence MLLLLARCFLVILASSLLVCPGLA ^ [C] GPGRGFG. [C] indicates the C24 position where palmitate is attached in the modified protein. Experiments relied on palmitoylation by endogenous expression of the Hedgehog acyltransferase HHAT, which was highly expressed (475 FPKM) in the parent clone of NIH3T3 cells. To achieve C-terminal cholesterol modification, the mouse ShhC domain was translationally fused to the C-terminus of the indicated synthetic protein, begging with the sequence - G[G]^CFPG… where [G] indicates the amino acid that will ultimately be modified with cholesterol in the processed proteoform. Nb23 was ordered as synthetic DNA based on the previously published sequence (*33*).

### Fast single particle tracking (fast SPT)

Sender cells were plated with receiver cells at a ratio of ~1:100 at ~90% confluence, i.e. 450,000 cells in 24-well glass-bottom plates (Cellvis). After one day when cells reached confluence, sender cells were induced with 50nM 4-OHT and allowed to secrete ligand overnight. Ligands were labelled with 50nM JF549i for 15 minutes at room temperature, and cells were washed 3 times with fresh media. Single particle microscopy was performed using TIRF microscopy as described previously(*13, 51*), with an exposure time of 6ms, and an imaging frequency of approximately 167Hz.

Single particle trajectories were measured using Quot, and the maximum likelihood diffusion rate of each molecule was estimated using SASPT(*24*). Exact parameters used by Quot and SASPT are reported in Supplemental table S1. Trajectories were not allowed to blink, and particles were allowed to jump 1.1µm per frame for fast SPT analysis.

Data were visualized using Python and ggplot2. To plot diffusion histograms, the state array from SASPT was marginalized across all localization errors and the argmax diffusion rate was calculated for each particle. The histogram bars reflect the log_10_ density of molecules at each diffusion rate. To plot the line, the state array from SASPT was marginalized across all localization errors and particles to generate the posterior probability density function for particles in each condition. The density traces were scaled for plotting on the same axis as the underlying maximum likelihood diffusion rate histograms.

To estimate the occupancy of each assigned molecular population, molecules were binned into diffusion rate intervals [0,0.1), [0.1,0.35), [0.35,2), [2,100] µm^2^/s, corresponding to molecules that were assigned categories “matrix,” “receptor,” “membrane,” or “free.”

To measure the bleaching process for single molecules, we extracted trajectories that were longer than 8 detections and sampled up to 250 trajectories per condition. For each trajectory, we re-calculated the raw intensity of the 3×3 pixel box centered on the reported spot position, and we estimated the background by calculating the median pixel intensity in a 7×7 pixel box with the same center. We performed this calculation for the entire trajectory, and for 10 frames after the trajectory ended at the last reported particle position. At every time point, we subtracted the background from the signal, and then for every trajectory we anchored the pre-bleaching value to 1, by dividing by the average signal value in the 10 frames before bleaching. After standardizing the intensity traces, we smoothed each trace over a 3-frame interval and plotted gray lines for individual traces. Next, we over-plotted a black line thar reflects the median adjusted signal at each time point. The same bleaching analysis approach was used for fast SPT and for slow SPT.

### Slow single particle tracking (slow SPT)

Imaging was performed as for fast SPT, except under minimal laser power and with 2500ms exposure time (corresponding to 1/4 Hz). Additionally, we used an iLas2 “ring TIRF” system (Gataca) to minimize illumination differences over the field of view. Molecules were tracked using Quot, allowing 150nm movement between consecutive frames. The data were filtered to obtain trajectories with at least 2 detections, and survival curves were plotted using Python and Matplotlib.

### Immunoprecipitation

SHH-Halo alleles were tagged with the HA tag, and Halo or Nb23-Halo alleles were tagged with the V5 tag. HA-tagged alleles were precipitated from conditioned media using anti-HA magnetic beads (Pierce #88836), and V5-tagged alleles were precipitated from conditioned media using Anti-V5 magnetic beads (Fisher #NC0777490). Precipitates were washed 3 times in phosphate-buffered saline (PBS) with 0.05% tween-20, then once in PBS. Protein eluted in 100 µg/mL recombinant HA or V5 peptide at 37°C for 10 minutes.

### CRISPR

Mutant cell lines were generated using CRISPR/Cas9. Briefly, plasmids expressing Cas9, mCherry and a target-specific sgRNA were transiently electroporated into wild-type NIH3T3 cells. Cells were sorted for mCherry expression by FACS two days after transfection, then recovered for 3-5 days. Cells were then re-sorted for single cells that had lost mCherry expression, and a separate pool of cells was collected to check bulk editing efficiency. Clones were ultimately sequenced using Plasmidsaurus amplicon sequencing, and mutation frequencies were calculated by computing the frequency of mutated k-mers at the sgRNA target site in the raw Plasmidsaurus Fastq files. sgRNA and screening primer sequences are listed in table S2. *Ptch1*^(*-*)^ cell lines were confirmed based on Sanger sequencing and were confirmed phenotypically based on loss of Nb23 receptor-binding activity. Cell lines generated in this study are listed in table S3.

### Specific signaling activity calculations

Conditioned media was treated with 50nM JF549i for >30 minutes to label Halo tag alleles to completion. Conditioned media was then analyzed using poly-acrylamide gel electrophoresis (PAGE), and gels were imaged using a Typhoon (GE Healthcare) fluorescence scanner. Because Halo-JF549i labeling is stoichiometric, we used the fluorescence of the bands in the PAGE gel to quantify the relative concentration of each signaling ligand in its respective conditioned media. Measurements were made using Fiji/ImageJ. To test the potency of each ligand, receiver cells were pre-grown to confluence, and then the media was replaced with media containing a concentration gradient of conditioned media. For diluted samples, the media was supplemented with conditioned media from wild-type NIH3T3 cells, such that across all ligand concentrations there was an equal amount of total conditioned media. Signaling response was measured after 48hr using flow cytometry, and the fraction of cells that were mCitrine-high were plotted using R/ggplot2.

### Gradient assays

NIH3T3 mouse fibroblasts were used as sender and receiver cells as described previously(*20*). Briefly, parental cell lines were modified with PiggyBAC to encode the transcriptional activator Gal4-ERT2 and a payload ligand as required for the experiment to generate 4-OHT-inducible sender cells. Receiver cells were modified using PiggyBAC to insert H2B-mCitrine under control of an 8x-Gli binding site array. Upon stimulation with Hedgehog, receiver cells induced the H2B-mCtirine reporter ~10-50-fold.

Sender cells were plated in a soft, bio-compatible 2-well culture insert (Ibidi) placed inside each well of a 6-well plate near confluence (50,000 cells in 50µL) and allowed to adhere to the cell culture substrate overnight. Then the 2-well culture insert was removed, and the well was over-plated with a confluent lawn of NIH3T3-derived Hedgehog-sensitive receiver cells, cGS401. This created a co-culture in which one zone included a mixture of sender and receiver cells, and the remainder of the well was confluently cultured with receiver cells only. After receiver cells adhered to the culture dish, sender cells were induced and allowed to signal for 36-48h and then imaged on a Nikon Ti2E epifluorescence microscope to measure the position of ligand senders (mCherry channel) and the intensity of the signaling response (mCitrine channel). Within an experiment, all images used the same illumination intensity, exposure settings, magnification and LUT scaling for plotting. Images were processed using Fiji/ImageJ

## Acknowledgements

We would like to thank Domenic Narducci, Matteo Mazzoco, Xavier Darzaq, Luke Lavis, Alex Cao, and Whitehead FACS facility. This work was supported by National Institute of Health grants DP2HD108777 (PL), R00HD087532 (PL), DP2GM140938 (ASH), R33CA257878 (ASH), UM1HG011536 (ASH), 1K99GM151487 (GS), Allen Distinguished Investigator Award, a Paul G. Allen Frontiers Group advised grant of the Paul G. Allen Family Foundation (PL), National Science Foundation grant 2036037 (ASH), and Jane Coffin Childs Fund Postdoctoral Fellowship (GS).

## Author contributions

GS and PL conceptualized the project. GS performed all the experiments and data analysis, with material support from ASH. The manuscript was written by GS and PL, with input from ASH.

## Data Sharing Plans

Plasmids used custom reagents used in this study are available upon request. Correspondence and requests for materials should be addressed to Pulin Li.

## Supplementary Materials

**Figure S1.**
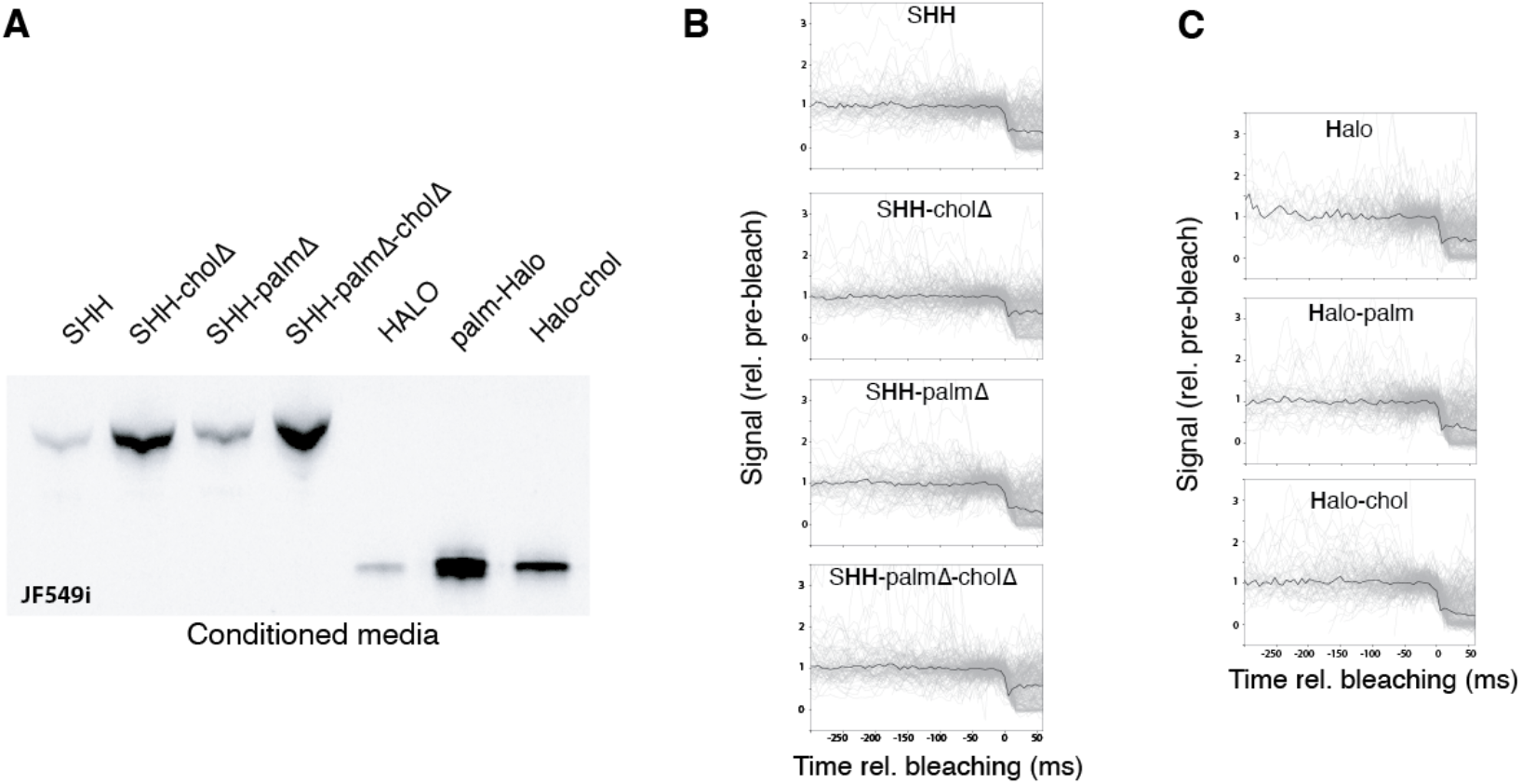
HALO-tagged proteins were successfully processed and were monomeric. A. Poly-acrylamide gel of conditioned media containing Hedgehog or Halo alleles labelled with JF549i. B. Bleaching curves for Hedgehog alleles calculated from fast SPT experiments. Gray lines represent individual bleaching traces, and black lines represent the median signal at each time step. C. Bleaching curves as in (B) for Halo alleles calculated from fast SPT.

**Figure S2.**
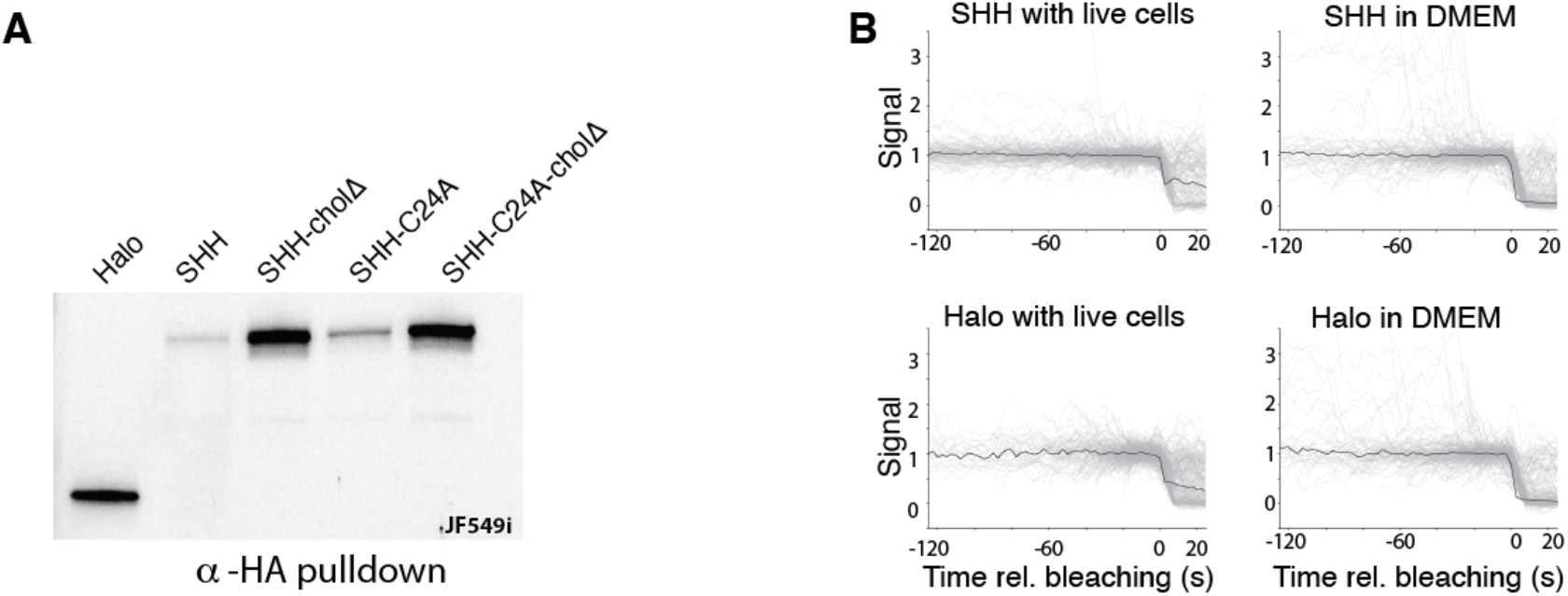
Purified proteins were monomeric in cell culture and in DMEM. A. Poly-acrylamide gel of immunoprecipitated Hedgehog and free Halo labelled with JF549i. B. Bleaching curves for purified Hedgehog alleles calculated from slow SPT experiments. Gray lines represent individual bleaching traces, and black lines represent the median signal at each time step.

**Figure S3.**
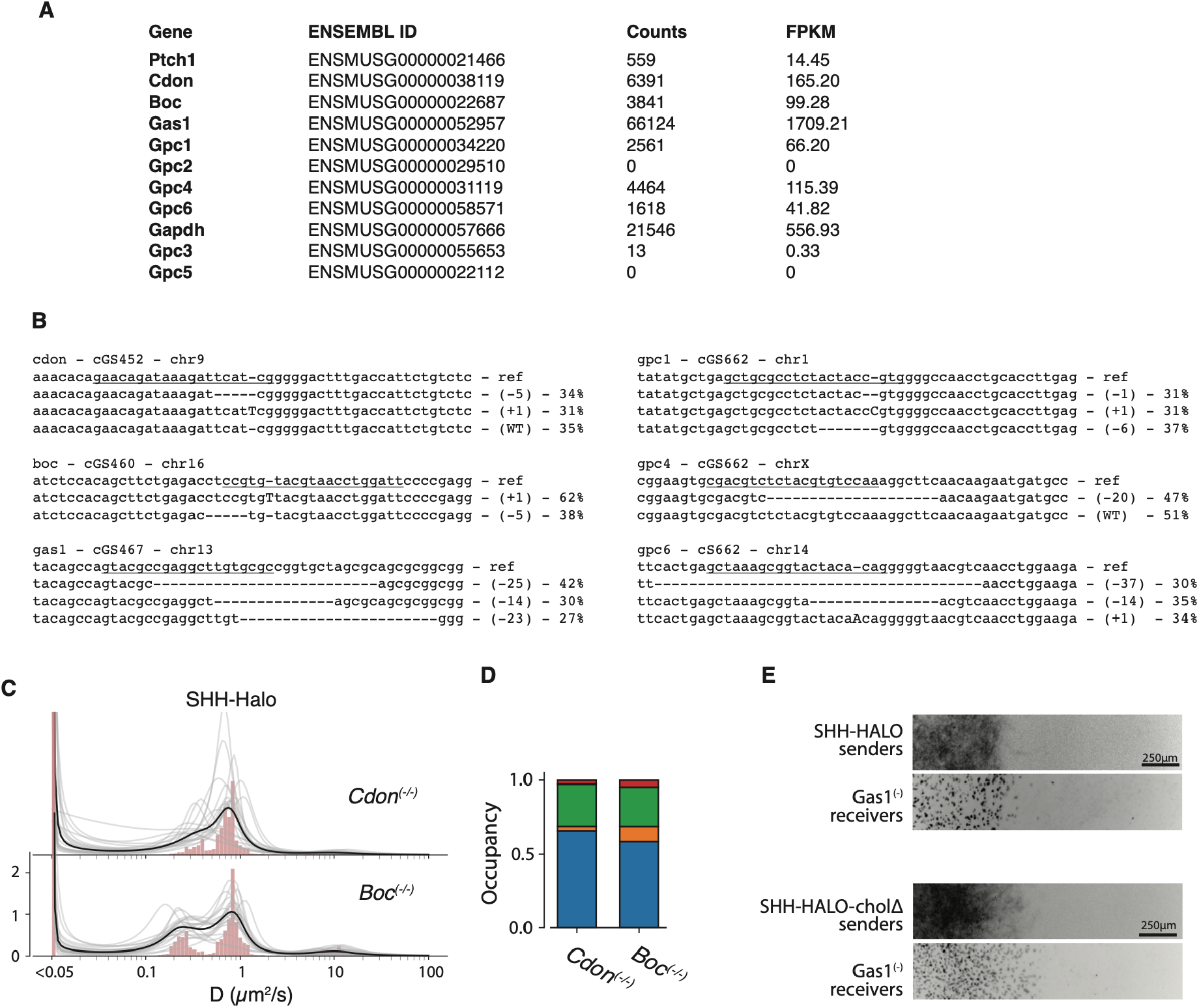
Genetic analysis of Hedgehog co-receptors. A. Expression level of Hedgehog co-receptors measured by RNAseq in wild-type NIH3T3 cells. B. Genotyping results for mutant NIH3T3-derived cell lines. Cell lines were triploid, consistent with prior observations about NIH3T3 cells. C. Histogram of diffusion rates for wild-type SHH diffusing among MEF-derived receiver cells of the indicated diploid genotypes. D. Quantification of state occupancy based on fast SPT in (C). E. Signaling gradients formed by SHH-derived ligands in Gas1^(-)^ receiver cells.

**Figure S4.**
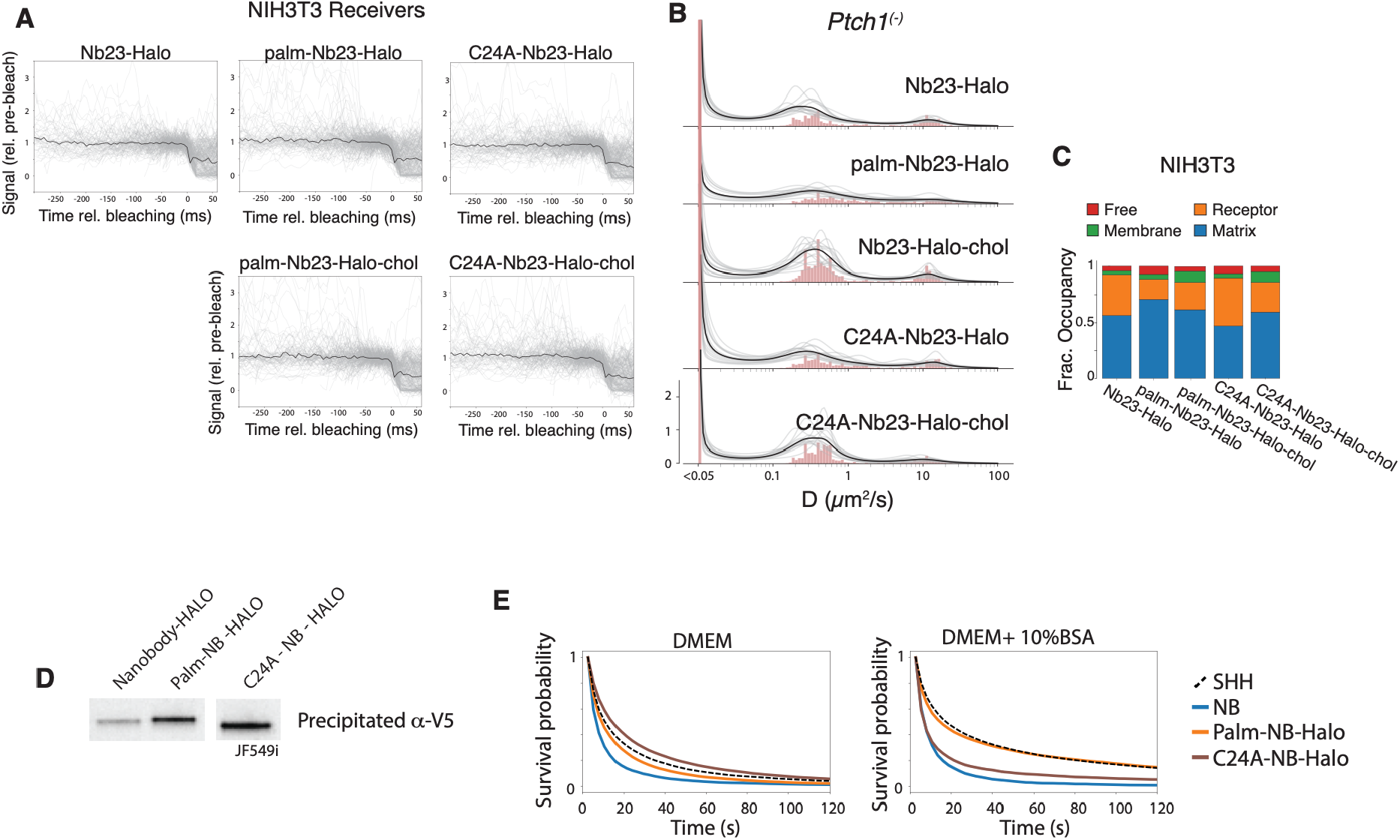
Nanobody alleles diffused as monomers in at Ptch1-dependent manner. A. Bleaching curves for Nb23 alleles calculated from fast SPT in Fig 4A. Gray lines represent individual bleaching traces, and black lines represent the median signal at each time step. B. Histogram of diffusion rates of Nb23 alleles diffusing among Ptch1^(-)^ receiver cells. C. Quantification of state occupancy in Fig 4A. D. Purification of JF549i-labeled nanobody alleles, visualized with JF549i. E. SlowSPT for purified nanobody-derived synthetic morphogens observed in the absence of receiver cells.

**Figure S5.**
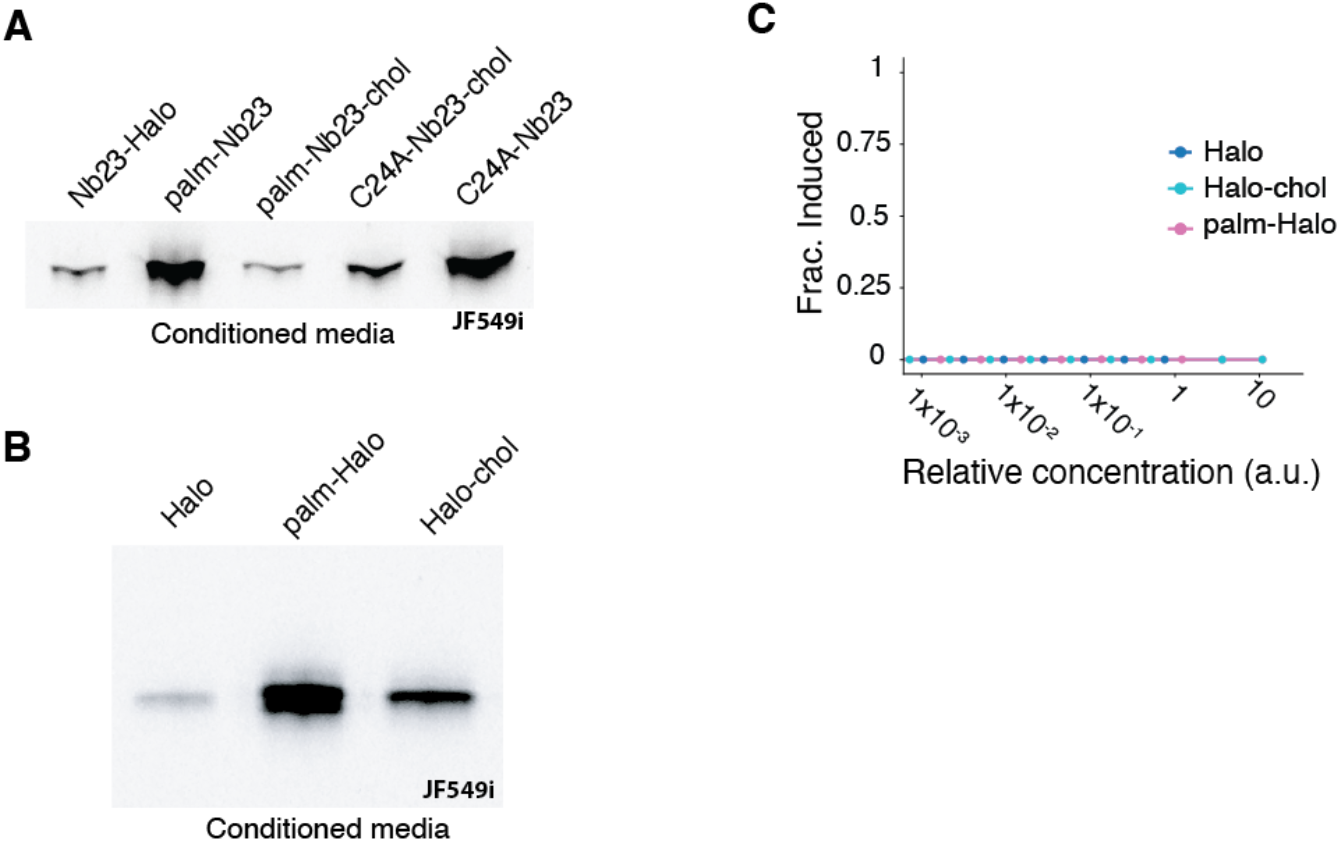
Halo-tagged allele expression and signaling activity. A. Conditioned media containing Nb23-Halo alleles labelled with JF549i, used to quantify the specific signaling activity of each Nb23-Halo allele. B. Conditioned media containing Halo alleles labelled with JF549i. C. Specific signaling activity of the Halo protein modified with N-terminal palmitate or C-terminal cholesterol, with NIH3T3-derived Hedgehog-sensitive reporter cells cGS401.

**Table S1.** Analysis parameters used by Quot and SASPT.

|  |  |
| --- | --- |
| quot ::<br>filter ::<br>start :: 0<br>method :: identity<br>chunk_size :: 100<br>detect ::<br>method :: llr<br>k :: 2.0<br>w :: 15<br>t :: 16.0<br>localize ::<br>method :: ls_int_gaussian<br>window_size :: 15<br>sigma :: 2.5<br>ridge :: 1e-05<br>max_iter :: 30<br>damp :: 1<br>camera_bg :: 108.0<br>track ::<br>method :: euclidean<br>pixel_size_um :: 0.11<br>frame_interval :: (determined from the data)<br>search_radius :: 1.1<br>max_blinks :: 0<br>min_I0 :: 100.0<br>scale :: 1.0 | saspt ::<br>likelihood_type :: rbme<br>pixel_size_um :: 0.11<br>frame_interval :: (determined from the data)<br>focal_depth :: 0.1<br>progress_bar :: False<br>splitsize :: 5<br>sample_size :: 100000<br>num_workers :: 12<br>diff_coefs :: np.power(10,(np.linspace(-2,2,125)))<br>loc_errors :: np.linspace(0.025 , 0.028 , 5)<br>start_frame :: 0 |

**Table S2.**
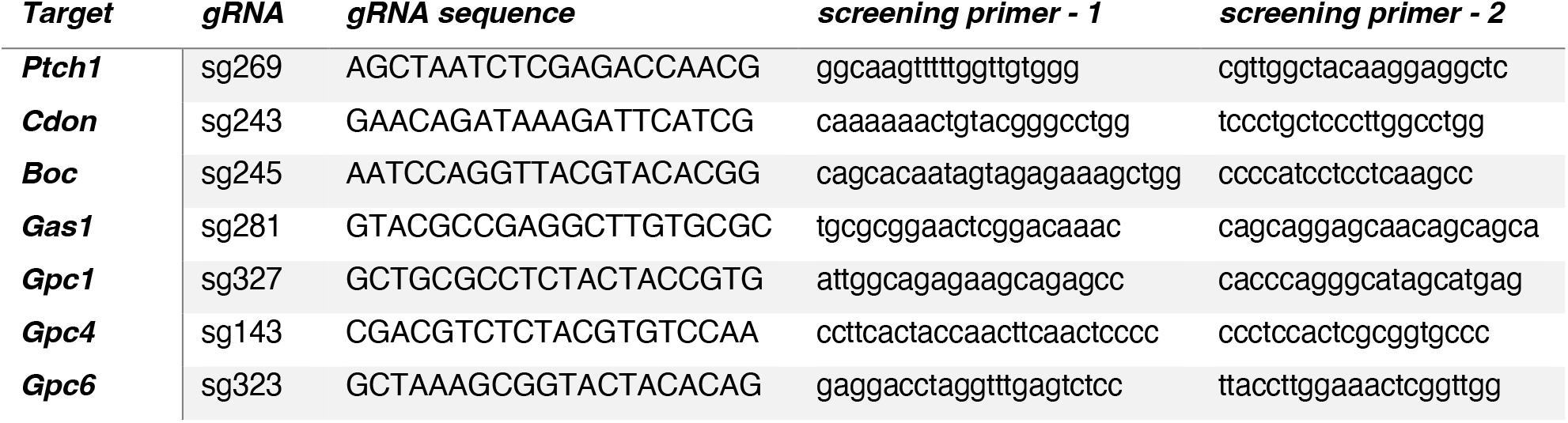
sgRNA sequences and screening primers for CRISPR mutagenesis of NIH3T3 cells.

**Table S3.** Clonal immortal cell lines generated and used in this study.

| *Cell line* | *genotype* | *confirmation method* | *notes* |
| --- | --- | --- | --- |
| **cGS572** | NIH3T3 |  | likely triploid, XXY |
| **cGS517** | NIH3T3 *ptch1*(-) | Sanger, phenotypic |  |
| **cGS452** | NIH3T3 *Cdon*(-) | Plasmidsaurus | (+0/-4/+1) |
| **cGS460** | NIH3T3 *Boc*(-) | Plasmidsaurus | (+1/+1/-5) |
| **cGS467** | NIH3T3 *Gas1*(-) | Plasmidsaurus | (-25/-14/-23) |
| **cGS662** | NIH3T3 *Gpc1*(-)<br>*Gpc4*(-) *Gpc6*(-) | Plasmidsaurus | *Gpc1*(-1/+1/-6) ; *Gpc4*(+0/-20) ;<br>*Gpc6*(-37/-14/+1) |

